# *Myelin regulatory factor* (*Myrf*) is a critical early regulator of retinal pigment epithelial development

**DOI:** 10.1101/2024.04.26.591281

**Authors:** Michelle L. Brinkmeier, Su Qing Wang, Hannah Pittman, Leonard Y. Cheung, Lev Prasov

**Author notes:** Author for correspondence: Lev Prasov, MD, PhD KEC 229 1000 Wall St. Ann Arbor, MI 48105.

## Abstract

Myelin regulatory factor (Myrf) is a critical transcription factor in early retinal and retinal pigment epithelial development, and human variants in *MYRF* are a cause for nanophthalmos. Single cell RNA sequencing (scRNAseq) was performed on *Myrf* conditional knockout mice (*Rx>Cre Myrf^fl/fl^*) at 3 developmental timepoints. *Myrf* was expressed specifically in the RPE, and expression was abrogated in *Rx>Cre Myrf^fl/fl^* eyes. scRNAseq analysis revealed a loss of RPE cells at all timepoints resulting from cell death. GO-term analysis in the RPE revealed downregulation of melanogenesis and anatomic structure morphogenesis pathways, which were supported by electron microscopy and histologic analysis. Novel structural target genes including *Ermn* and *Upk3b*, along with macular degeneration and inherited retinal disease genes were identified as downregulated, and a strong upregulation of TGFß/BMP signaling and effectors was observed. Regulon analysis placed *Myrf* downstream of *Pax6* and *Mitf* and upstream of *Sox10* in RPE differentiation. Together, these results suggest a strong role for Myrf in the RPE maturation by regulating melanogenesis, cell survival, and cell structure, in part acting through suppression of TGFß signaling and activation of *Sox10*.

**SUMMARY STATEMENT:** *Myrf* regulates RPE development, melanogenesis, and is important for cell structure and survival, in part through regulation of *Ermn*, *Upk3b* and *Sox10,* and BMP/TGFb signaling.

## INTRODUCTION

The retinal pigmented epithelium, RPE, is a single layer of polarized epithelium that develops between the neural retina and the choroidal vasculature and sclera (Marmorstein et al., 1998). The RPE provides support critical for maintenance of the outer segments of the photoreceptors and regulates transport of nutrients and waste between the choroid vasculature and the neural retina. The RPE is formed from the dorsal outer layer of the optic vesicle, which is derived from the anterior neural plate under the direction of eye field transcription factors including *Lhx2, Nr2e1, Otx2, Rax, Six3, Six6, Tbx3,* and *Pax6* (Amram et al., 2017; Diacou et al., 2022). *Pax6* together with *Pax2* and *Otx2* regulate the expression of the earliest marker of the pigmented RPE, *Mitf.* In turn, *Mitf* is a master regulator of RPE development. Loss of *Mitf* leads to transdifferentiation of RPE to neural retina and ectopic expression of *Mitf* activates genes involved in melanogenesis (Horsford et al., 2005; Nguyen and Arnheiter, 2000).

Many signaling pathways are important in the differentiation, maintenance, and function of the RPE (reviewed in (Amram et al., 2017) and (Cardozo et al., 2020)). TGFß, BMP, and WNT ligands in the periocular mesenchyme and surface ectoderm surrounding the lens drive the RPE fate (Steinfeld et al., 2013). Inactivation of WNT signaling in the early forming RPE inhibits the differentiation of RPE, likely through ß*-catenin* activation of *Otx2* and *Mitf* (Fujimura et al., 2009; Westenskow et al., 2009)*. Bmp7* is expressed throughout the RPE and loss of *Bmp7* in a mouse model leads to failure of eye development (Morcillo et al., 2006). *Bmp4* is expressed in the dorsal region of the developing eye and loss of *Bmp4* leads to transdifferentiation of neural retina to RPE (Huang et al., 2015). In addition, *Bmp2* in the RPE has been shown to be a negative regulator of eye growth in chick studies (Sakuta et al., 2006; Zhang et al., 2012). In mice, postnatal eye growth is regulated by repression of *Lrp2* and *Bmp2* by *Srebp2* (Mai et al., 2022). The Hippo pathway also regulates RPE development and specification. Loss of YAP in early optic cup stages of eye development results in a transdifferentiation of RPE to retina (Kim et al., 2016). Conditional deletion of YAP with the *Rx>cre* results in hypopigmentation of the RPE, a folding and thinning of the retinal layer, transdifferentiation of the RPE into retina, and altered expression of key RPE transcription factors. YAP is also crucial in maintaining the polarity of the RPE which impacts the survival of the retinal progenitor cells (Kim et al., 2016).

Maintenance of structural integrity and cellular polarity in RPE is critical to its function and the survival of overlying photoreceptors (reviewed in (Eamegdool et al., 2020) and (Gelat et al., 2022)). Adherens and tight junctions formed between the epithelial cells create a barrier between the vasculature of the choroid and the retinal cells while allowing for the transport of metabolites, nutrients, and the phagocytosis and disposal of photoreceptor outer segments. In addition, the basement membrane of the RPE cells sits on Bruch’s membrane which provides support to the RPE cells and strengthens the choroid/retinal barrier. Epithelial to mesenchymal transition (EMT) of the RPE cells occurs in many ocular disease states including proliferative vitreoretinopathy (PVR) and age-related macular degeneration (AMD). TGFß signaling, specifically through TGFB2, has been identified as a driving mechanism of EMT leading to compromised RPE structure and disease progression (Hachana and Larrivee, 2022).

*Myelin regulatory factor* (*MYRF*) is a membrane-associated transcription factor critical for development and maintenance of myelination. Pathogenic MYRF variants have been identified in families with autosomal dominantly inherited isolated nanophthalmos, a condition characterized by a small, structurally normal eye, as well as a feature of ocular cardiac urogenital syndrome (Calonga-Solis et al., 2022; Guo et al., 2019; Gupta et al., 2022; Hagedorn et al., 2020; Kaplan et al., 1993; Qi et al., 2018; Siggs et al., 2020; Siggs et al., 2019). We previously showed that biallelic loss of *Myrf* in the developing eye results in dysfunction of the retinal pigmented epithelium and subsequent retinal degeneration (Garnai et al., 2019; Yu et al., 2021b). However, the precise site of action and function of *Myrf* in eye development has been debated (Yu et al., 2021a; Yu et al., 2021b). To elucidate the role of *Myrf* in eye development and place it in the hierarchy of RPE differentiation, we performed single cell RNAseq (scRNAseq) on early eye field conditional knockout *Myrf* mouse model, *Rx>cre Myrf fl/fl,* at multiple developmental time points. Our findings place *Myrf* within the hierarchy of transcription factors known to regulate RPE development and help define the molecular mechanisms and signaling pathways driving RPE dysfunction in the absence of *Myrf*.

## RESULTS

### MYRF is specifically expressed in the retinal pigment epithelium

While dominant mutations in MYRF have been definitively shown to cause nanophthalmos in humans, the specific cellular drivers of phenotype remain a subject of debate. There is speculation that the ciliary margin/ciliary zonules (Yu et al., 2021a), RPE (Garnai et al., 2019), and retina (Ouyang et al., 2022) may all be implicated. To better define specific cell-type of action for MYRF, we systemically evaluated the expression pattern of *Myrf* in mouse using RNAscope *in situ* hybridization. We found that *Myrf* mRNA is predominantly expressed in the RPE throughout retinal development (Figure 1 A-D), with much lower or no expression in the ciliary body (Figure 1K) and retina. Conditional deletion of *Myrf* in the early eye field using an *Rx>Cre* transgene leads to loss of *Myrf* mRNA signal in the RPE, supporting the specificity of the probes (Figure 1 E-H). Optic nerve expression of *Myrf* is consistent with its known role in regulation of myelination, and is not altered by *Rx>cre* deletion, given that developing oligodendrocytes or progenitors do not express *Rx* (Figure 1 I-J). These data are consistent with prior qRT-PCR data from the human eye (Garnai et al., 2019) and supported by existing RNAseq datasets (Figure 2B). Together, these data support the fact that the RPE is the primary site of function for MYRF within the eye.

**Figure 1.**
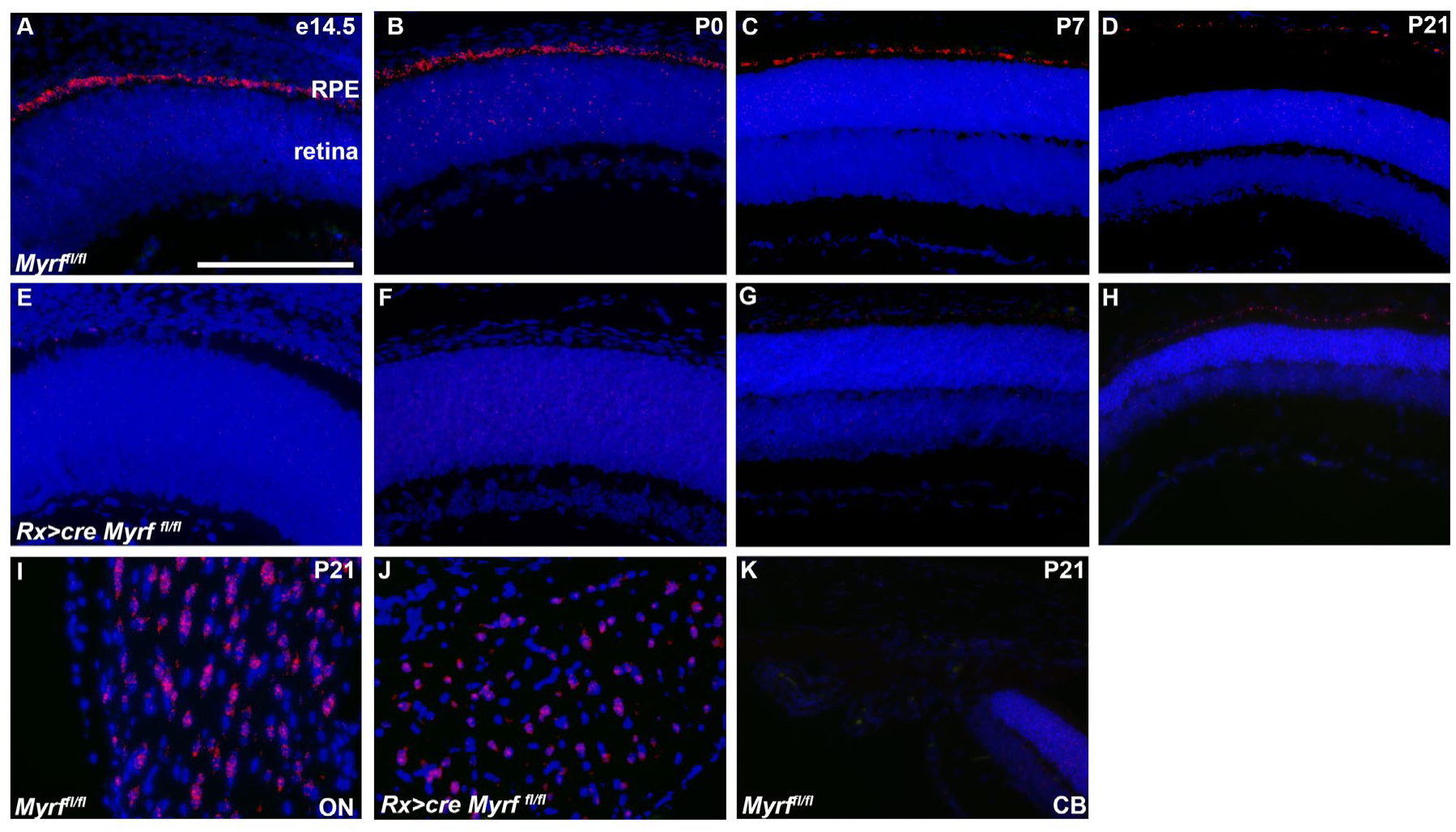
*Myrf* is specifically expressed in the RPE during mouse development. (A-H) *Myrf* transcripts in the RPE were detected by RNAscope during mouse embryonic development through postnatal day 21 in *Myrf^fl/fl^* controls (A-D) and absent in *Rx>cre Myrf^fl/fl^* mutants (E-H). (I-J) Expression of *Myrf* in the optic nerve (ON) serves as a positive control (I-J) and was not abrogated with *Rx>Cre* deletion. (K) Expression of *Myrf* is not detected in the ciliary body (CB, K). Scale bar, 200uM.

**Figure 2.**
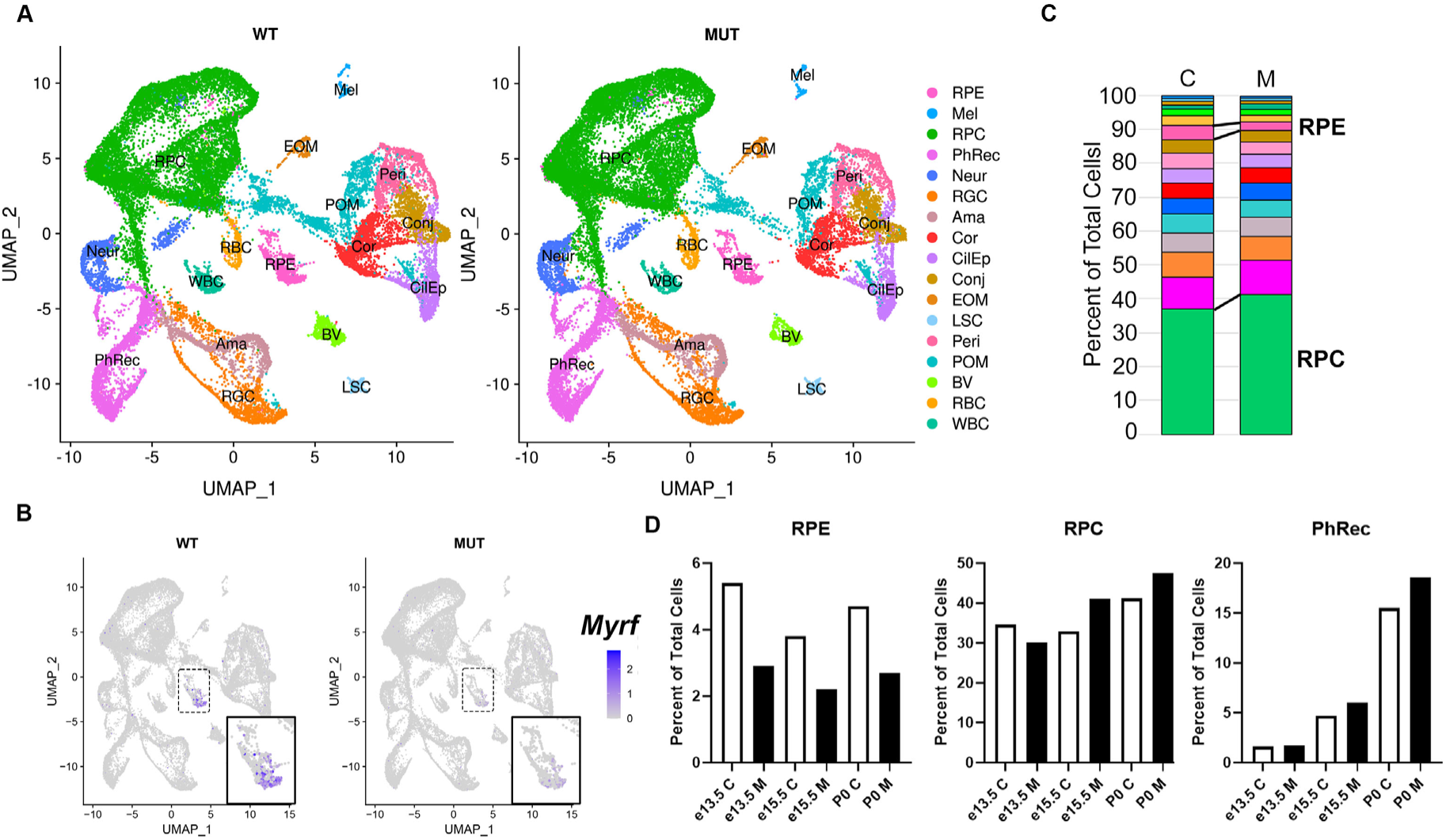
**scRNAseq analysis *Rx>*c*re Myrf^fl/fl^* mice suggests cell population shifts in the RPE and RPCs**. (A) Combined UMAP plots from e13.5, e15.5, and P0 *Myrf^fl/fl^* (n=3 pooled for each time point) and *Rx>*c*re Myrf^fl/fl^* (n=3 pooled for each time point) samples showing cell populations and cluster identification. (B) UMAP plot showing expression of *Myrf* specific to the RPE cluster in the control group and absent in the mutant group. (C) Stacked bar chart showing population distribution of clusters as a percentage of the total number of cells (control, C; mutant, M). The RPE cluster is highlighted as being decreased in the mutant and the RPC as being increased. (D) Bar charts show the decrease in cell number within the RPE mutant (M) cluster across all time points, an increase in the RPC cluster during later time points, and no change in the photoreceptor cluster (PhRec).

### Loss of MYRF alters cell type distributions during development

To define the mechanism by which loss of MYRF leads to retinal degeneration (Garnai et al., 2019) and eye size defects, we performed scRNAseq using the 10X genomics platform to characterize the gene expression changes within the RPE and neighboring cell types in *Rx>*c*re Myrf^fl/fl^* mice as compared to *Myrf^fl/fl^* controls using 3 specific developmental timepoints: embryonic day (e)13.5 – before histologic phenotypes are apparent in the *Rx>*c*re Myrf^fl/fl^* eyes (Garnai et al., 2019); e15.5 – the first time at which overt RPE pigmentation phenotypes appear; and postnatal day (P)0 – a time at which secondary changes may be apparent in neighboring cell types. We collected 6039, 13017, and 14305 cells for e13.5, e15.5, and P0 wildtype, respectively and 6734, 13880, and 12222, cells for e13.5, e15.5, and P0 mutant timepoints respectively. The median genes per cell were comparable in wild-type and mutant samples (2826, 2985, and 1915 for e13.5, e15.5, and P0 *Myrf^fl/fl^* and 2957, 2862, and 2179 for respective *RxCre Myrf^fl/fl^* eyes. After quality control filtering for dead cells, doublets, and poor-quality cells and combination and integration of the data, we conducted unsupervised clustering and were able to define 28 clusters, with 17 showing distinct gene expression patterns (Figure 2A, Supplemental Figure 1). The 7 retinal progenitor clusters, 3 photoreceptor clusters, 3 retinal ganglion cell clusters, and 2 periocular mesenchyme clusters that showed similar gene expression patterns and position on the UMAP dimensionality reduction plot were thus combined into the respective RPC, PhRec, RGC, and POM clusters for further analysis. Analysis of *Myrf* transcripts within the clusters again suggested specific expression within the RPE during development, and loss of expression in *Rx>Cre Myrf^fl/fl^* eye cups as expected (Figure 2B). All cell types were represented in both the *Rx>*c*re Myrf^fl/fl^* eyes and controls, but there were population differences with a reduction in RPE cell proportions in the mutant mice across all three developmental time points (5% vs. 3% at e13.5, 4% vs.2% at e15.5MUT, 5% vs. 3% at P0) and an increase in RPC numbers at e15.5 and P0 (33% vs. 41% at e15.5, 41% vs. 48% at P0). No changes were observed in the photoreceptor clusters across timepoints or genotype (Figure 2C, D), suggesting photoreceptor loss occurs later than P0.

### MYRF loss leads to RPE cell death without altered cell cycle dynamics

To further define the source of population level differences, we systematically examined cell cycle dynamics and cell death among the RPE and RPC populations. Cleaved caspase-3 staining revealed an increase in cell death at e15.5, while there was no cell death in *Myrf^fl/fl^* or *Rx>*c*re Myrf^fl/fl^* controls at postnatal (P0, P3) timepoints (Supplemental Figure 2). To further quantify cell death, we systematically evaluated apoptosis in the RPE at e14.5 using Terminal deoxynucleotidyl transferase dUTP nick end labeling (TUNEL), a timepoint immediately prior to the observable gross phenotype of RPE depigmentation. Co-staining with OTX2 was used to definitively mark RPE cells and identify retinal progenitors (low OTX2 in the neuroblastic layer) as well as nascent developing photoreceptors (high OTX2, basolateral) (Figure 3A-C). Apoptotic cells were identified in the RPE of both *Rx>cre Myrf^+/fl^* and *Rx>*c*re Myrf^fl/fl^* mice, but very rarely in *Myrf^fl/fl^* controls (insets, Figure 3A’-C’). The percentage of TUNEL positive RPE cells was found to be significantly different in *Myrf^fl/fl^* mice and Rx>cre *Myrf^fl/fl^* eyes (0.6±0.7% vs. 11±9%, p = 0.01), and between *Myrf^fl/fl^* and *Rx>*c*re Myrf^fl/+^* eyes (0.6±0.7% vs. 5±5%, p =0.03) (Figure 3D). To exclude altered cellular proliferation and cell cycle dynamics in RPE accounting for population differences, we conducted cell cycle analysis in Seurat, which demonstrated similar levels of G1, S, and G2/M phase cells across the genotypes at each time point (Supplemental Figure 3). Subclustering of RPE followed by pseudotime trajectory analysis using Monocle3 reveals a modest decrease in cells with higher pseudotime values in the P0 timepoint, suggesting a differentiation delay in the P0 *Rx>cre Myrf^fl/fl^* RPE (Supplemental Figure 4). The number of OTX2 expressing retinal progenitors (OTX2 low) and OTX2 expressing developing photoreceptors (OTX2 high) was unchanged between genotypes (Figure 3D) confirming our scRNA sequencing findings in photoreceptor cell population. These studies confirm that the RPE cell population changes in *Rx>cre Myrf^fl/fl^* eyes are due to apoptosis, and that photoreceptor loss occurs after P0.

**Figure 3.**
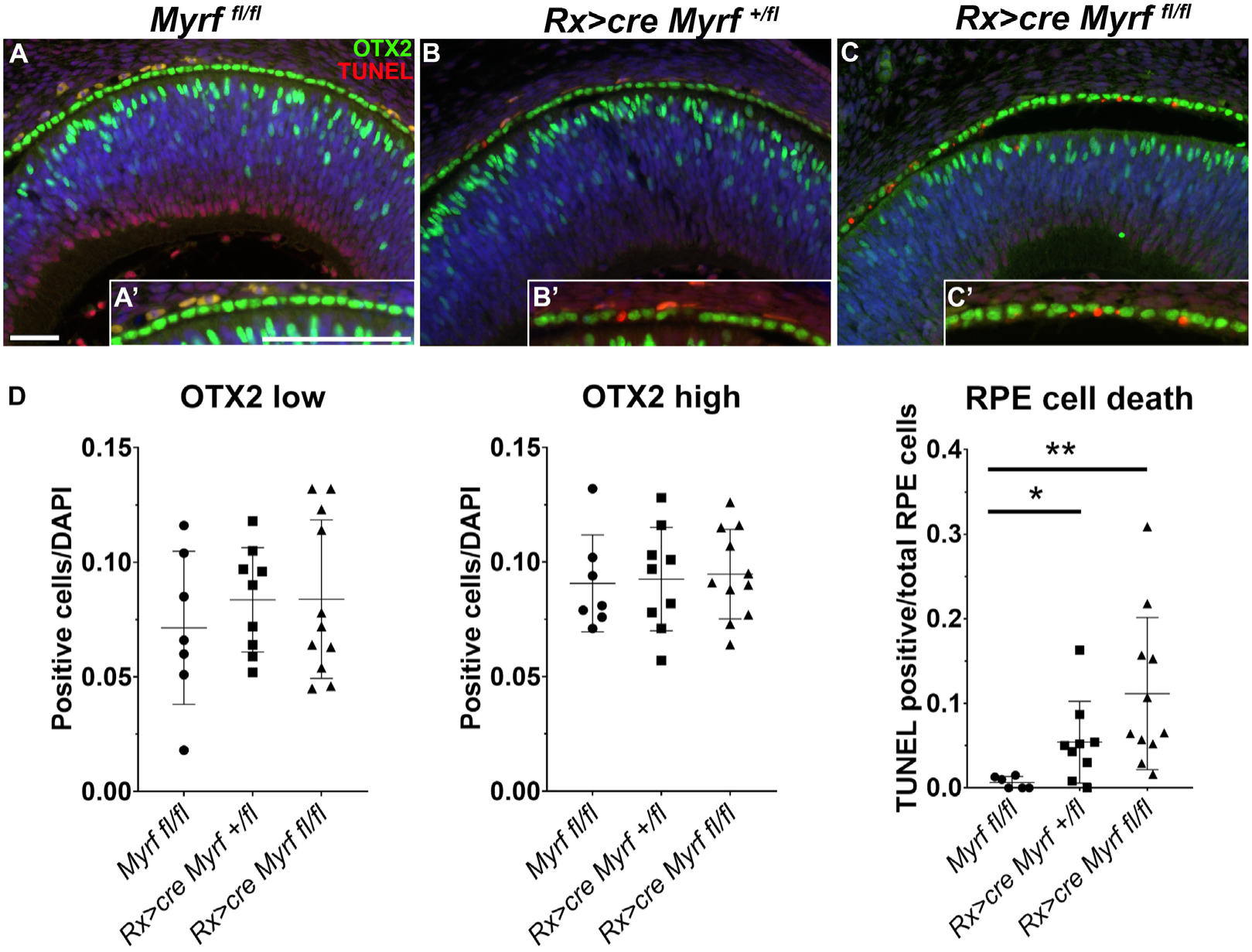
RPE cell death occurs in *Rx>*c*re Myrf^fl/fl^* mutants. Apoptotic cells were identified by TUNEL staining (red) in *Myrf^fl/fl^*, n=7(A), *Rx>*c*re Myrf^+/fl^*, n=9 (B), and *Rx>cre Myrf^fl/fl^,* n=10(C) mice at embryonic day e14.5. OTX2 expression (green) was used to mark the RPE, retinal progenitor cells (OTX2 low) and developing photoreceptor cells (OTX2 high). Higher magnification insets were used to demonstrate the increase in TUNEL positive cells in the *Rx>cre Myrf^fl/fl^* RPE compared to controls (A’-C’). D. TUNEL positive cells within the RPE monolayer (OTX2 positive) were quantitated and compared to the total number of DAPI+ cells within the RPE. Unpaired Student’s T Test was used for statistical analysis, **=p=0.01, *=p<0.05 Scale bar, 50 um.

### *Myrf* loss alters conserved biological pathways in RPE development

To define genes regulated by *Myrf* and key cellular pathways responsible for the RPE development and maturation, we conducted differentially expressed genes analysis (DEGs) within the RPE cluster at each developmental timepoint. Loss of *Myrf* is observed at all timepoints and reduction of *Tmem98* can be seen starting at e15.5, consistent with our prior report (Garnai et al., 2019). We identified many genes known to be associated with inherited retinal disease (IRD) and age-related macular degeneration (AMD) that are downregulated in the absence of *Myrf.* These include *Unc119* and *Rlbp1* starting at e13.5 and *Cdh3, Arhgef18, Ctsd, Rgr, Rdh5, Timp3, Slc16a8, Col8a1,* and *Cfh* at P0, suggesting that *Myrf* plays a key role in regulating RPE disease associated genes (Figure 4A). To further explore pathways impacted by loss of *Myrf* and define novel gene candidates for RPE disease, we analyzed enrichment of Gene Ontology (GO terms) on differentially expressed transcripts from the RPE cluster using the using the PANTHER Overrepresentation Test (Release 20200728) and the GO Ontology database DOI: 10.5281/zenodo.3954044 Released 2020-07-16. GO terms significantly enriched in genes that are downregulated in the *Rx>*c*re Myrf^fl/fl^* mutant include regulation of melanin biosynthetic process (MelBio, 75 fold, p = 0.005), visual perception (VisPer, 11 fold, p = 0.05), extracellular matrix organization (ECM, 8 fold, p = 0.02), organic hydroxy compound metabolic process (HydCompMet, 6 fold, p = 0.01), blood vessel development (BV, 5 fold, p = 0.02), tube morphogenesis (TM, 5 fold, p = 0.02), regulation of anatomical structure morphogenesis (AnaStrMor, 4 fold, p = 0.02), and positive regulation of cellular process (RegCellPro, 2 fold, p = 0.03) (Figure 4B). GO terms enriched in genes that are upregulated in the *Rx>Cre Myrf^fl/fl^* mutant include positive regulation of cellular proliferation (PosRegCellProlif, 8-fold, p = 0.02) and cellular response to chemical stimulus (CellRespChemStim, 6-fold, p = 0.01) (Figure 4C). Within these GO terms, underlying genes that were differentially expressed across all three timepoints were defined to compile candidates that are more likely to be direct targets of *Myrf* (Figure 4D) from the complete list of DEGs (Supplemental Table 1). Critical regulators of melanogenesis (*Pmel, Slc24a5, Tyrp1, Cdh2),* visual perception*(Lum, Cfh, Rdh10, Rdh5),* extracellular matrix (*Vit, Fbln2, Aplp1, Ramp2),* hydroxy compound metabolism *(Ttr, Rbp1, Dct),* blood vessels *(Bmpt, Thbs1, Slc1l1, Fzd4),* tube morphogenesis *(Sox10, Mgp, Lrp1),* anatomical structure *(Ezr, F3, Kif1a, Ermn),* and regulation of cellular process (*Acsk1, Cst3, Otx2, Trf*) are downregulated. Key regulators of cellular proliferation and cellular respiration (*Enpp2, Gas6, Ptn, Ctgf, Id1, Id3, Fgf15, Igfbp2*) are upregulated.

**Figure 4.**
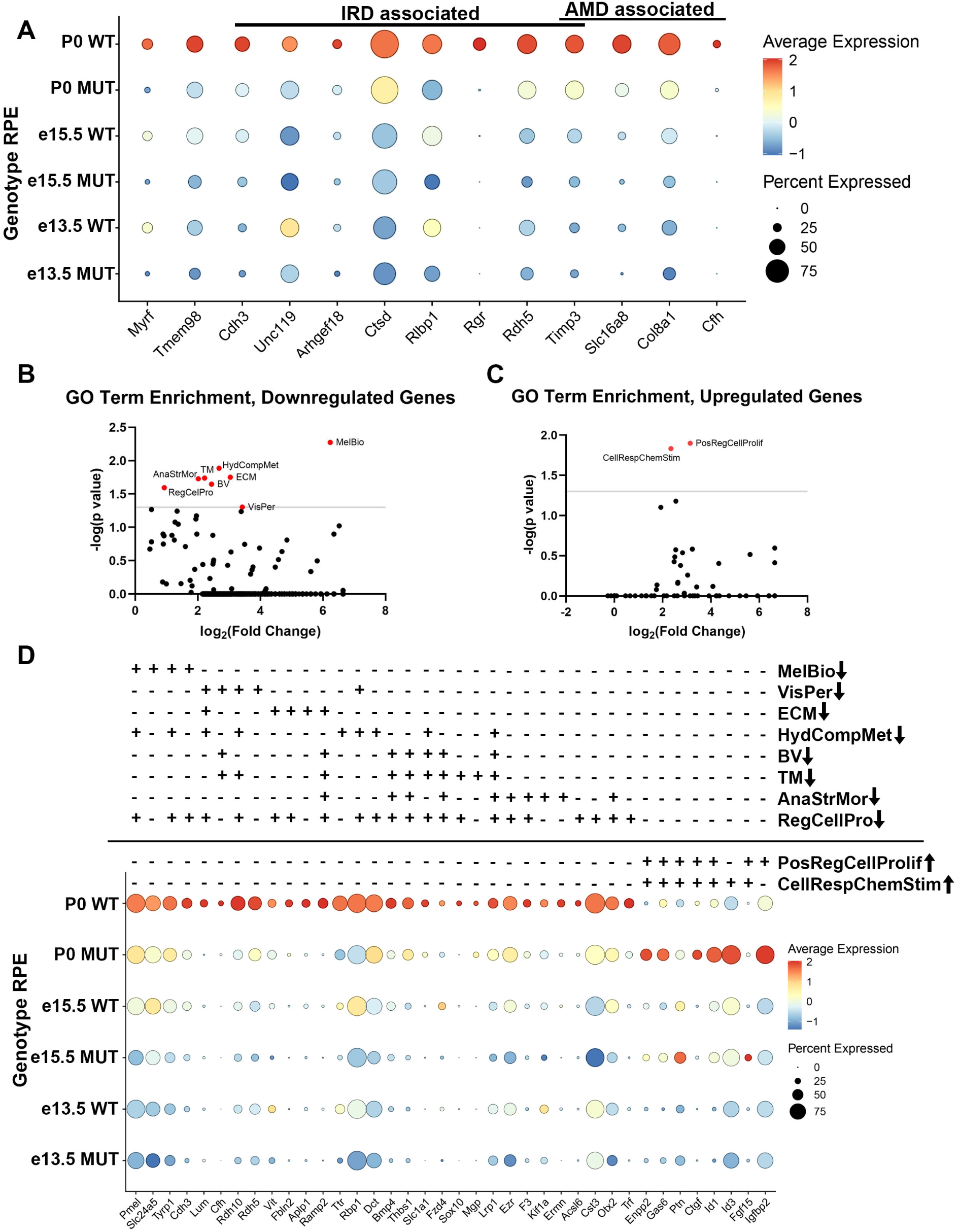
Loss of *Myrf* during development alters critical pathways in RPE development and disease. (A) DotPlot analysis showing the impact of loss of *Myrf* on expression of genes known to play a role in Inherited Retinal Diseases (IRD) and Age-related Macular Degeneration (AMD). (B-C) PANTHER GO terms enriched in downregulated (B) or upregulated (C) genes in the RPE cluster differential expression analysis of *Rx>Cre Myrf^fl/fl^* vs. *Myrf^fl/fl^* controls across all timepoints. (D) DotPlot analysis of selected genes from enriched GO terms across biological pathways, highlighting changes in genes involved in regulation of melanin biosynthetic process (MelBio), visual perception (VisPer), extracellular matrix organization (ECM), organic hydroxy compound metabolic process (HydCompMet), blood vessel development (BV), tube morphogenesis (TM), regulation of anatomical structure morphogenesis (AnaStrMor), positive regulation of cellular proliferation (PosRegCellProlif), cellular response to chemical stimulus (CellRespChemStim).

### MYRF regulates key cytoskeletal proteins and alters RPE morphology

Enrichment of melanin biosynthetic process and extracellular matrix organization suggests *Rx>*c*re Myrf^fl/fl^* mutants may have structural changes in the RPE and are consistent with the observed loss of pigmentation in *Rx>*c*re Myrf^fl/fl^* mice. In principle, loss of pigmentation could be related either to loss of melanosomes or reduced melanin. To define ultrastructural features and the source of depigmentation in *Myrf* loss-of-function, we conducted Transmission Electron Microscopy (TEM) in *Myrf^fl/fl^* and *Rx>*c*re Myrf^fl/fl^* mice at P21 (Figure 5). *Rx>*c*re Myrf^fl/fl^* mutants exhibit abnormal apical microvilli structure and organization, loss of photoreceptors, abnormal outer segment (OS) structure, and thickening of Bruch’s membrane (Figure 5A-B). Additionally, *Rx>*c*re Myrf^fl/fl^* displayed a significant reduction in the number of melanosomes in the RPE (4.9± 2.3×10^5^/mm^2^ vs. 1.6±0.7×10^5^/mm^2^, p=0.0103), suggesting loss of melanosomes underly the depigmentation phenotype in *Myrf* loss-of-function mice (Figure 5A-C). Consistent with changes in ultrastructural organization, *Rx>*c*re Myrf^fl/fl^* mutants had reduced expression of the apically localized cytoskeletal protein *Ermn* (Liang et al., 2018) and extracellular matrix marker *Upk3b* (Figure 6). FeaturePlots from the scRNAseq data reveal specific expression of *Ermn* and *Upk3b* within the RPE cluster in *Myrf^fl/fl^* dataset and loss of expression in the *Rx>*c*re Myrf^fl/fl^* dataset (Figure 6A), with onset of *Ermn* expression at e13.5 and increasing expression through development and onset of *Upk3b* expression at P0. To validate dysregulation of ERMN expression in *Rx>*c*re Myrf^fl/fl^* eyes, immunohistochemical analysis was performed at P21. Co-localization of apical cytoskeletal marker ezrin (EZR) and ERMN were observed in *Myrf^fl/fl^* control RPE (Figure 6B), but ERMN expression was reduced in in *Rx>*c*re Myrf^fl/fl^* mutants, while EZR retained apical localization (Figure 6C). Specificity of expression of extracellular matrix gene, *Upk3b*, to the RPE and its loss in *Rx>*c*re Myrf^fl/f^*^l^ eyes at P0 were confirmed by RNA scope *in situ* hybridization in the absence of a validated antibody (Figure 6D). The structural changes observed in the *Rx>cre Myrf^fl/fl^* mutants and the loss of ERMN and *Upk3b* expression support a role for MYRF in RPE structure and microvilli organization.

**Figure 5.**
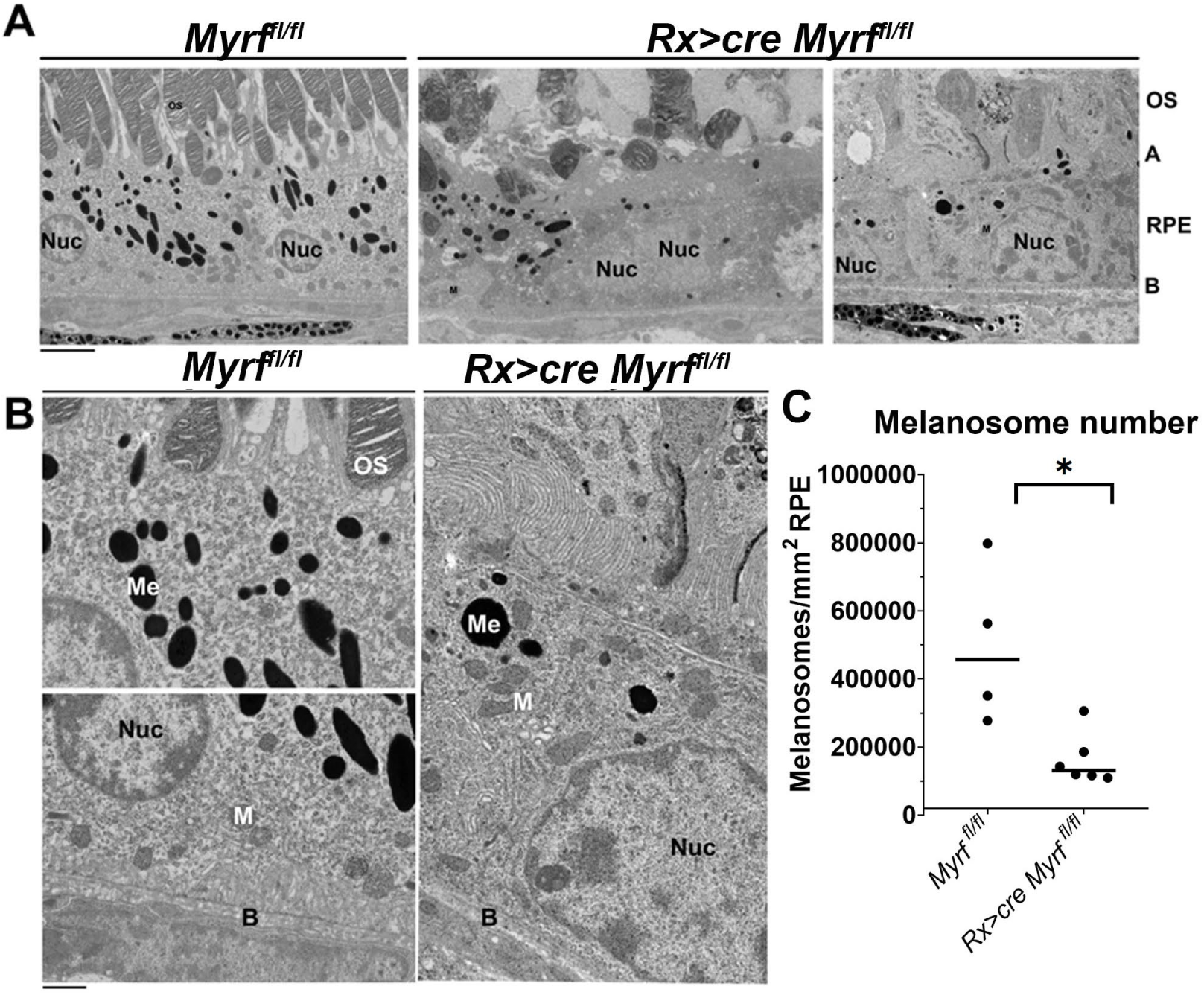
Changes in ultrastructural features in *Rx>*c*re Myrf^fl/fl^* RPE identified in TEM. Low (A) and high (B) magnification transmission electron micrograph (TEM) images were generated to analyze the ultrastructure, and number of melanosomes (Me), Bruch’s membrane (B), apical microvillous structure in the RPE and outer segment (OS) structure and in *Myrf^fl/fl^* (n=4) controls and *Rx>cre Myrf^fl/fl^* (n=6) mutants at P21. C. Quantification of the number of melanosomes per mm^2^ RPE. Scale bar in A, 4um; in B, 1um. A, apical surface; M, mitochondria; *, p<0.05.

**Figure 6.**
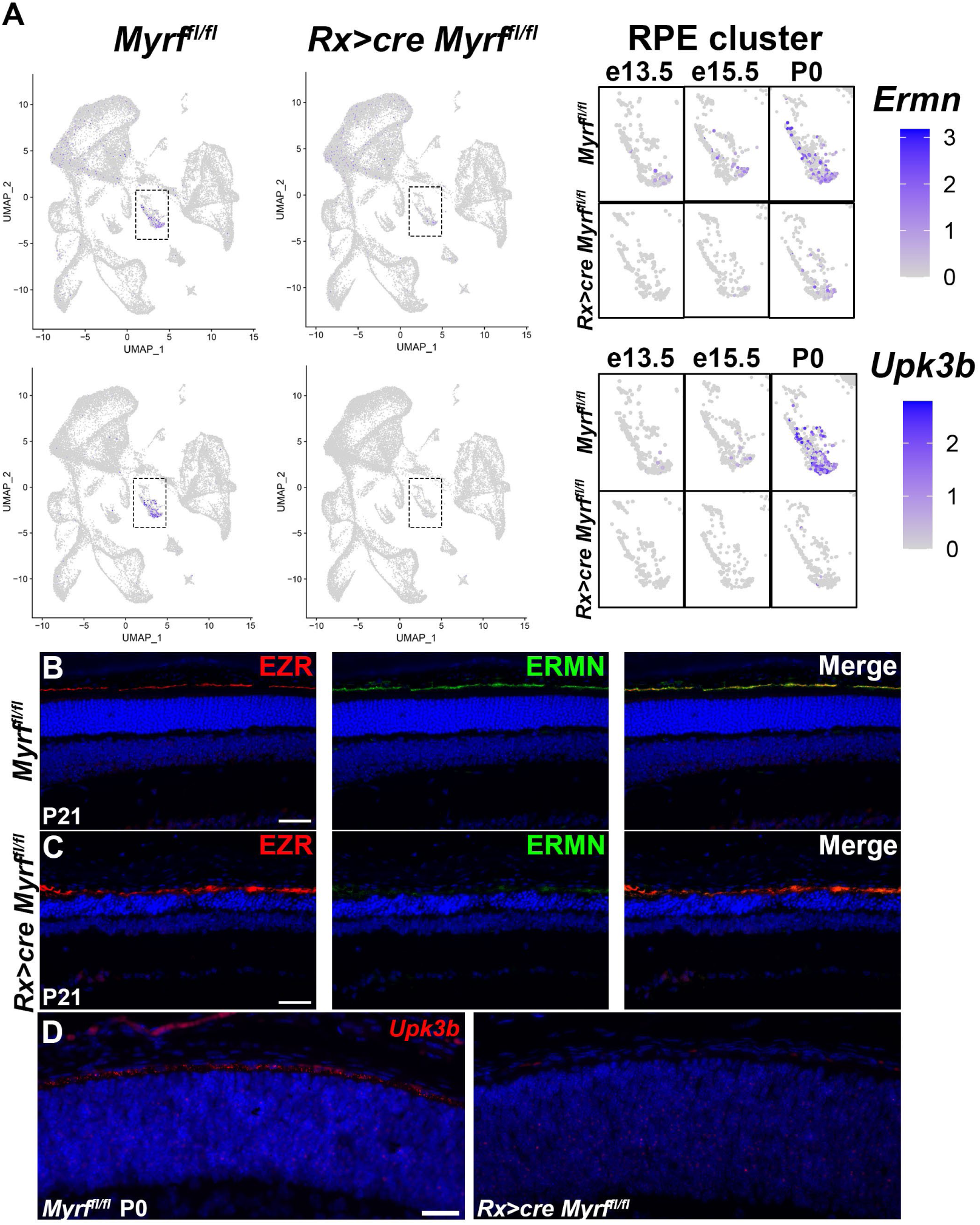
MYRF regulates expression of cytoskeletal and extracellular matrix proteins. (A) FeaturePlot analysis of expression of *Ermn* and *Upk3b* showing specificity of expression to the RPE cluster and loss of expression in the *Rx>cre Myrf^fl/fl^* mutants. Larger images show expression within the RPE cluster through development. (B, C) Cytoskeletal proteins were marked within the RPE using antibodies to Ezrin (EZR, red) and Ermin (ERMN, green). (B) P21 sections of *Myrf^fl/fl^* control eyes (n=2) show both EZR and ERMN localized to the RPE. (C) P21 sections of *Rx>cre Myrf^fl/fl^* mutants (n=3) show retention of EZR expression and loss of ERMN expression within the RPE. (D) RNAscope of *Upk3b* at P0 shows expression specific to the RPE layer and loss of expression in the *Rx>cre Myrf^fl/fl^* mutants (n=3) compared to *Myrf^fl/fl^* controls (n=4). Scale bars in all panels represent 50um.

### Loss of MYRF leads to altered transcription factor regulation including loss of SOX10 expression in RPE

To define the place of MYRF in the regulatory hierarchy of RPE development, we conducted Single-Cell rEgulatory Network Inference and Clustering (SCENIC) analysis (Aibar et al., 2017; Van de Sande et al., 2020). SCENIC is a computational tool that analyzes the co-expression of transcription factors with other genes in the dataset and considering putative cis-regulatory binding motifs present in the proposed target genes to define regulons. A higher regulon activity is assigned to transcription factors co-expressed with targets containing putative binding motifs upstream of the transcription start site (Figure 7A, Supplemental Table 2). Within the RPE cluster, transcription factors known to be important in early stages of RPE development including *Vax2* and *Pax6* have high regulon activity at e13.5 and e15.5 and those important later in development including *Otx2* and *Vsx2* have higher regulon activity at P0. *Mitf, Gsx2, Vsx2, Otx2* regulons were among those unaffected by *Myrf* loss through multiple RPE developmental timepoints, suggesting that *Myrf* does not directly regulate these genes or their targets. Consistent with this, we see no significant differences in transcription factor expression levels among these (Supplemental Figure 5). In contrast, we see a significant reduction in *Sox10* regulon activity within the RPE at P0. Given that SOX10 and MYRF have been identified as co-regulators of oligodendrocyte differentiation and maturation (Bernhardt et al., 2022; Sock and Wegner, 2021), we next investigated whether a similar regulatory network may exist in RPE*. Sox10* transcripts were found to be significantly reduced in the *Rx>*c*re Myrf^fl/fl^* mutant RPE DEG list and identified in two categories in the GO term enrichment analysis. We analyzed the distribution of *Sox10* transcripts across our scRNAseq clusters and found they were downregulated specifically in the *Rx>*c*re Myrf^fl/fl^* RPE cluster compared to *Myrf^fl/fl^* controls across all time points, but not changed in the melanocyte cluster (Figure 7B). Consistent with these results, SOX10 protein is expressed in both the melanocytes and RPE of *Myrf^fl/fl^* controls at e13.5, e15.5, and P3 (Figure 7C-E). While SOX10 expression is retained in the melanocytes of *Rx>*c*re Myrf^fl/fl^* mutants, it is absent from the RPE cells. These results suggest that *Sox10* expression is dependent on *Myrf* within the RPE.

**Figure 7.**
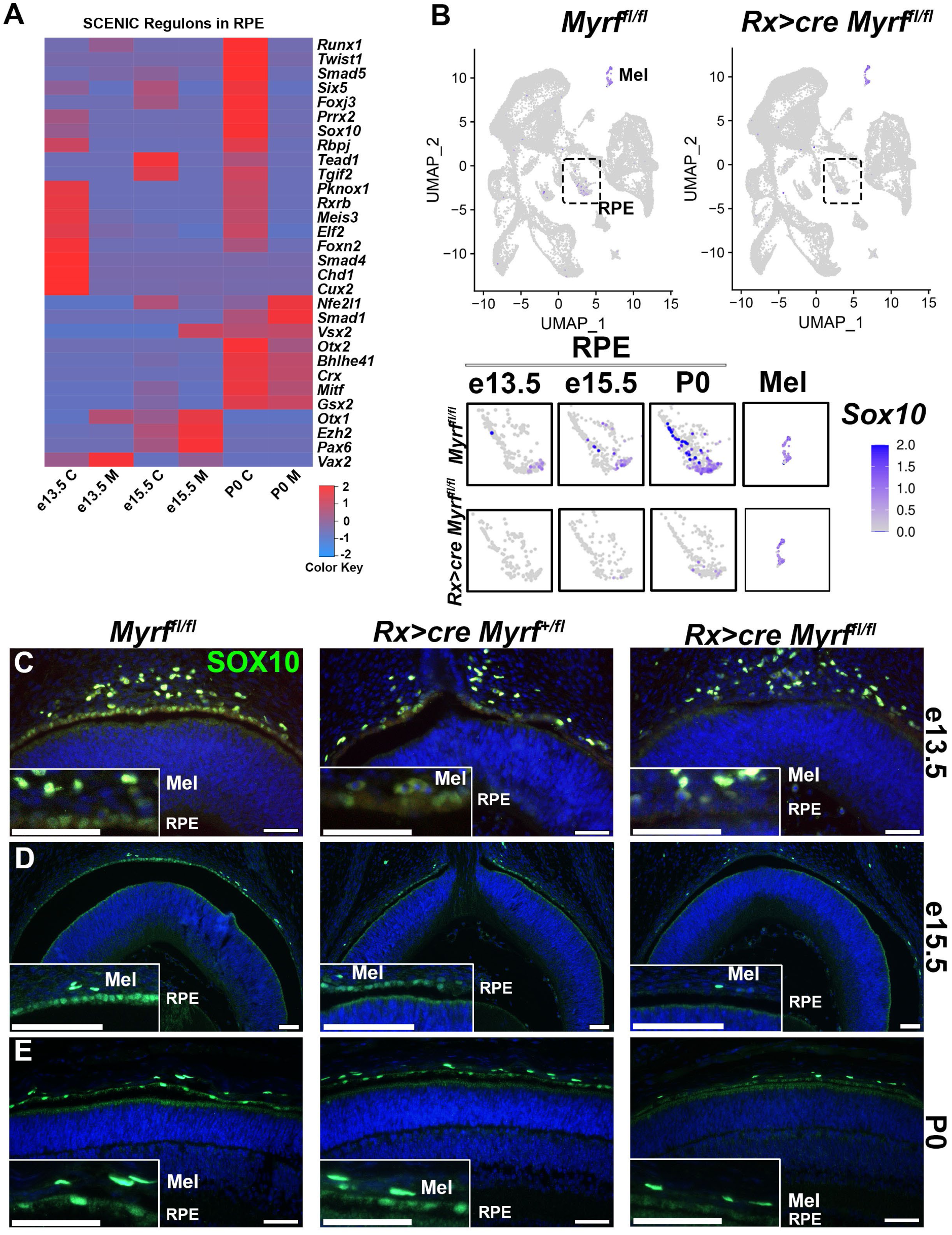
*Myrf* regulates SOX10 expression in RPE cells. (A) Single-Cell rEgulatory Network Inference and Clustering (SCENIC) analysis within the RPE cluster from the scRNAseq data infers active transcription factors across all developmental time points including members of the TGFB signaling pathway. (B) FeaturePlot analysis of scRNAseq transcripts shows *Sox10* expression in both the melanocyte and RPE clusters in *Myrf^fl/fl^* controls. Expression is retained in the melanocyte cluster in *Rx>cre Myrf^fl/fl^* mutants and lost in the RPE cluster. Larger images of the RPE cluster demonstrate onset of expression in the *Myrf^fl/fl^* controls at e13.5 and increasing through P0. Only a few cells containing *Sox10* transcripts can be identified in the *Rx>cre Myrf^fl/fl^* mutant at later time points. Larger image of the melanocyte cluster shows no changes in expression. (C-E) SOX10 protein is localized to the melanocytes and RPE in *Myrf^fl/fl^* and *Rx>cre Myrf^+/fl^* controls at e13.5 (*Myrf^fl/fl^* (n=3), *Rx>cre Myrf^+/fl^* (n=3), *Rx>cre Myrf^fl/fl^* (n=3)) (C), e15.5 (*Myrf^fl/fl^* (n=2), *Rx>cre Myrf^+/fl^* (n=3), *Rx>cre Myrf^fl/fl^* (n=3)) (D), and P3 (*Myrf^fl/fl^* (n=3), *Rx>cre Myrf^+/fl^* (n=2), *Rx>cre Myrf^fl/fl^* (n=3)) (E) SOX10 expression is only retained in the melanocytes of *Rx>cre Myrf^fl/fl^* mutants. Scale bars, 50um.

### MYRF regulates genes involved in proliferation, including BMP/TGFß signaling

Further investigation of regulon dysregulation in the RPE revealed many members of the BMP/TGFß signaling pathway, including *Smad5, Tgif2, Smad4,* and *Smad1* showing differential regulon activity between the *Myrf^fl/fl^* and *Rx>*c*re Myrf^fl/fl^* RPE clusters (Figure 7A). Given the role of BMP/TGFß signaling in cellular communication and differentiation, we used the computational tools MultiNichenetr (Browaeys et al., 2023) and Differential Nichenetr (Browaeys et al., 2020) to predict ligand, receptor, target gene interactions within the RPE cluster (Figure 8A-B, Supplemental Figure 6). Consistent with the SCENIC analysis, differential interactions within the BMP/TGFß signaling were also identified, including *Bmp4, Tgfb2,* and *Bmp2*. Interestingly, the expression and ligand activity of *Bmp2,* a known regulator of eye growth, is increased in the mutants, and multiple targets are also present within the RPE cluster. Expression analysis of BMP/TGFß signaling pathway members within the RPE cluster throughout development (Figure 8C) revealed dysregulation of TGFß/BMP components and effectors within the RPE. A known inhibitor of the pathway, *Wfikkn2*, and its downstream target, *Gdf11*, are decreased in the *Rx>*c*re Myrf^fl/fl^* RPE clusters. Consistent with the loss of inhibition, increases in expression of *Tgfb2, Bmp2, Smad1, Smad6, Smad9, Id1, Id2,* and *Id3* were also observed. Based on the Seurat VlnPlot, the expression of *Wfikkn2* is specific to the RPE cluster and lost in the *Rx>cre Myrf^fl/fl^* scRNAseq datasets (Figure 9A). We confirmed loss of *Wfikkn2* in the *Rx>cre Myrf^fl/fl^* mice at P0 using RNAscope (Figure 9B). Ligands of the TGFß pathway, *Tgfb2* and *Bmp2* were both elevated in the RPE cluster in our scRNAseq dataset, as seen with the VlnPlot, and our mouse model. (Figure 9A, C, D). TGFB2 is expressed in the RPC layer of *Myrf^fl/fl^* eyes at P0, with little to no expression in the RPE layer (Figure 9C). Elevated TGFB2 expression was observed in both the RPE and RPC layers of *Rx>cre Myrf^fl/fl^* mutants. Elevated levels of *Bmp2* transcripts were also detected in the RPE layer using *Bmp2* RNAscope (Figure 9D). Consistent with increases in BMP/TGFß signaling, the *Bmp2* inducible transcription factor *Id3*, showed increased expression within the RPE cluster of *Rx>*c*re Myrf fl/fl* mutants (Figure 9A), and this was confirmed by RNAscope (Figure 9E). Activation of BMP/TGFß signaling leads to phosphorylation of activating SMAD transcription factors (i.e. SMAD1/5/9). We investigated phosphorylated SMAD, pSMAD, as a marker of signaling activity. pSMAD 1/5/9 was not detected at P0 in *Myrf ^fl/fl^* controls (Figure 9F) but detected in the RPE of *Rx>cre Myrf^fl/fl^* mutants. Together our results support that loss of *Myrf* leads to activation of BMP/TGFß signaling and its effectors, linking *Myrf* to RPE dysfunction and/or eye growth pathways.

**Figure 8.**
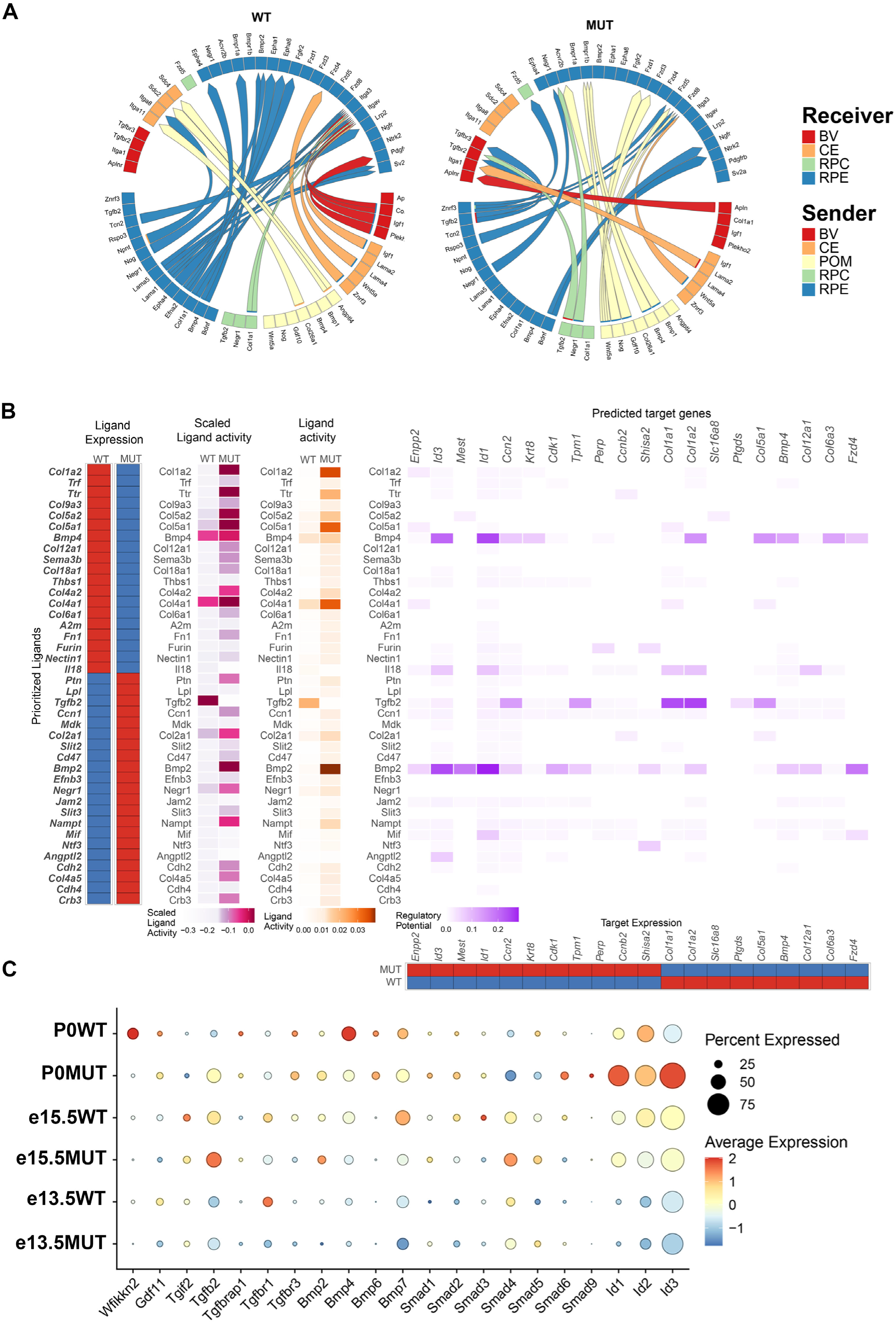
Computational analysis of the RPE cluster suggests alterations in the TGFB signaling pathway. (A) Circos plot generated from MultiNichenetr analysis predicting ligand receptor partners in control and mutant scRNAseq data sets from a combination of senders and receivers. (B) Analysis of changes in predicted signaling pathways within the RPE cluster using Differential Nichenetr. Expression and activity of ligands within the RPE is seen on the left and predicted targets within the RPE based on the Differential Nichenetr pathway analysis is seen on the right side. (C) DotPlot analysis of TGFB family members expressed in the RPE cluster across all genotypes highlights differential expression between *Myrf^fl/fl^* and *Rx>cre Myrf^fl/fl^* cells. (C-*Myrf^fl/fl^*; M-*Rx>cre Myrf^fl/fl^*)

**Figure 9.**
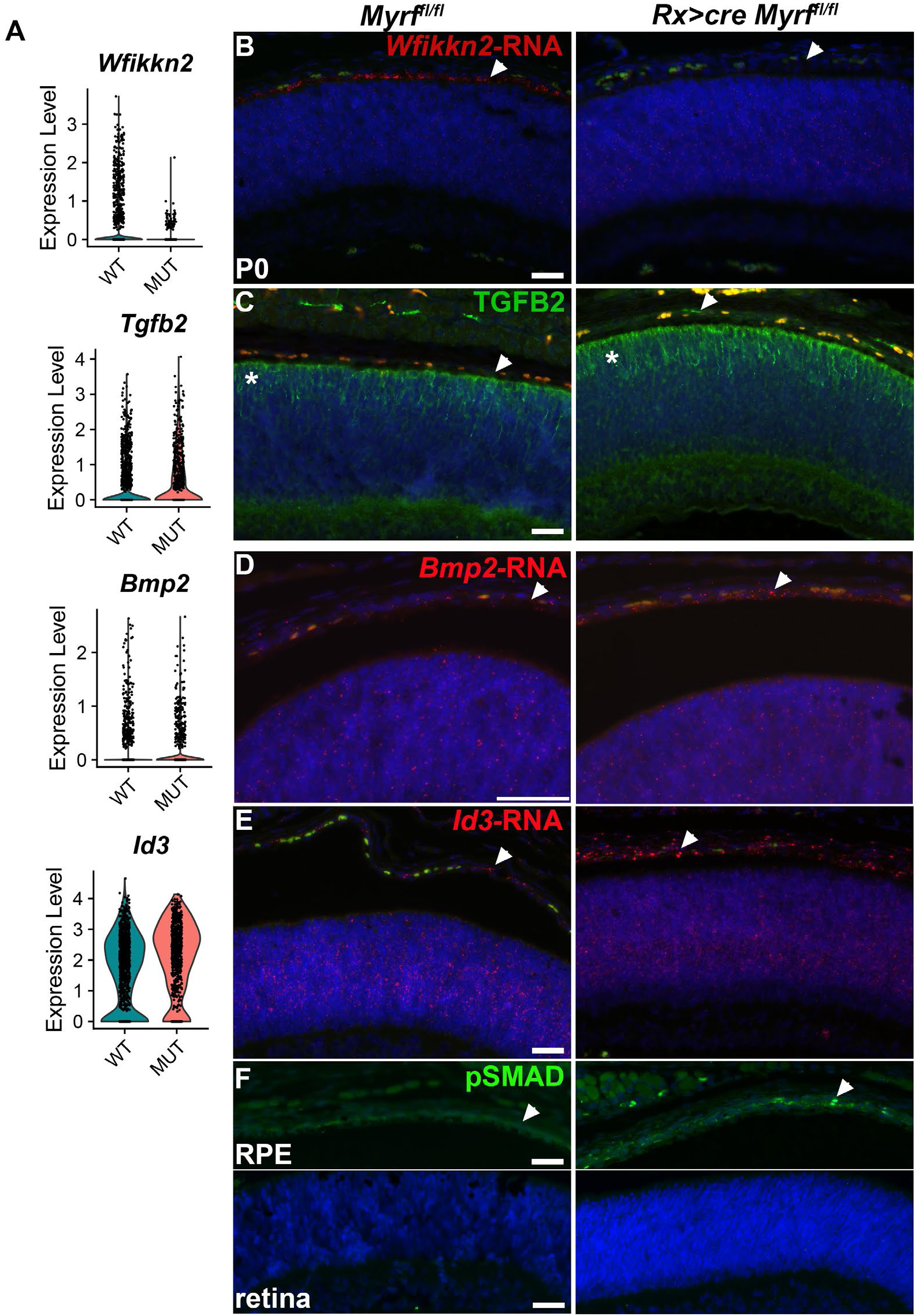
Increased BMP/TGFB signaling pathway activity in *Rx>cre Myrf^fl/fl^* mutants. (A) VlnPlot representation of transcript levels of *Wfikkn2*, *Tgfb2*, *Bmp2,* and *Id3* in the RPE cluster highlighting a decrease in the pathway inhibitor, *Wfikkn2* and increases in *Tgfb2, Bmp2,* and *Id3*. (B) RNAscope of *Wfikkn2* confirms specificity to RPE layer in *Myrf^fl/fl^* (n=4) and loss of expression in the *Rx>cre Myrf^fl/fl^* mutants (n=3) at P0. (C) TGFB2 protein is observed in the RPC of *Myrf^fl/fl^* controls (n=4) at P0, but not in the RPE layer. TGFB2 is present in the RPE layer of *Rx>cre Myrf^fl/fl^* mutants (n=3) and increased levels of expression are also seen in the RPC layer. (D) RNAscope of *Bmp2* demonstrates elevated expression in the RPE of *Rx>cre Myrf^fl/fl^* mutants (n=6) compared to *Myrf^fl/fl^* controls (n=4). (E) RNAscope of *Id3* shows increased expression in the RPE layer of *Rx>cre Myrf^fl/fl^* mutants (n=4) compared to controls (n=4). (F) Phosphorylated SMAD 1/5/9 immunostaining indicates an increase in activity within the RPE layer of *Rx>cre Myrf^fl/fl^* mutants (n=3) compared to no staining in *Myrf^fl/fl^* controls (n=3). (RPC-retinal progenitor cells; RPE-retinal pigmented epithelial cells; arrowheads-RPE layer, asterisk-RPC layer). Scale bars in all panels indicate 50um.

## DISCUSSION

The exact localization and function of *Myrf* during eye development is controversial. We have provided multiple lines of evidence supporting RPE predominant expression of *Myrf* during mouse ocular development, suggesting its primary role is within the RPE. Our initial studies with human *MYRF* expression demonstrated RNA specificity to the RPE and optic nerve, with no transcripts identified in the retina or ciliary body by qPCR (Garnai et al., 2019). We also confirmed in mouse that *Myrf* transcripts were over 100-fold more abundant in RPE as compared to retina (Garnai et al., 2019). In addition, we provided antibody staining in mice and human samples, depicting specific staining in the RPE and high non-specific background levels throughout the retina even in *Myrf* knockout mice. Compiled aggregated RNAseq data from the National Eye Institute EyeIntegration Pan Human Gene Expression plots support exponentially higher levels of *MYRF* expression in the RPE compared to the cornea or retina (Supplemental Figure 7). Our current data with RNAscope *in situ* hybridization in control and *Rx>cre Myrf^fl/fl^* mutants throughout development and postnatal time points supports that *Myrf* transcripts are specifically found in the RPE at least by E13.5 and show loss of expression in the *Myrf* conditional knockout mice. Further, our scRNAseq data detect *Myrf* transcripts only in the RPE cluster. The vast number of mouse models in which defects in the RPE lead to photoreceptor impairment (*Atg5* and *Atg7* (Zhang et al., 2017), *Ift20* (Kretschmer et al., 2023), *Mfrp* (Fogerty and Besharse, 2011), *Dapl1* (Ma et al., 2023), *Tsc1* (Go et al., 2020), *Kcnj13* (Roman et al., 2018), *Sod2* (Mao et al., 2014), *Rb1cc1* (Yao et al., 2015), *Lamp2* (Notomi et al., 2019), *Bsg* (Han et al., 2020), *Pten* (Kim et al., 2008), and *RPE65* (Gouras et al., 2002)) and the preservation of RPC progenitors and early photoreceptors in our *Rx>Cre Myrf^fl/fl^* mice support that phenotypes in these mice are primarily RPE-driven. Further studies with conditional deletion of *Myrf* in the retina and RPE separately will be needed to definitively demonstrate this.

Our results support that *Myrf* is vital for cell survival, melanogenesis, and establishing structural integrity of RPE. Other mouse models of reduced *Myrf* in eye development, including one model of a heterozygous patient variant and one heterozygous for a loss of function allele, suggest MYRF may play a role in the ciliary body and retina, respectively (Yu et al., 2021a; Yu et al., 2021b). Based on the age of mice analyzed, the phenotypes observed in these mouse models may be secondary to loss of MYRF in the RPE or a result of off target CRISPR effects. The genes altered in these mouse models are not different in our scRNAseq dataset (Supplemental Figure 8). In our model, while RPE specification is unaffected in *Rx>cre Myrf^fl/fl^* mice and the initial steps of differentiation appear unaltered with a normal appearing RPE monolayer at E13.5, we observe significant gene expression changes by scRNA sequencing even at these early time points. Apoptotic RPE cells are identified in the *Rx>cre Myrf^fl/fl^* mutants during embryonic development and an overall reduction in proportion of RPE cells in the eye cups is observed, preceding the loss of pigmentation. The RPE pigmentation defects appear to be driven by loss of melanosomes number and reductions in expression of melanosome associated genes, such as *Pmel* and *Tyrp1*.

Our ultrastructural and gene expression analysis reveals disruption and disorganization of the microvilli on the apical surface of the RPE cells, which leads to loss of their intimate contact with the photoreceptor outer segments. This interaction is critical to the function of the RPE cells in phagocytosis of photoreceptor outer segments and contributes to the degeneration of the retinal layer. A thickening of Bruch’s membrane can also be seen in the *Rx>cre Myrf^fl/fl^* mutants. A similar increase in thickness of Bruch’s membrane is also observed in *Lamp2* deficient mice as the RPE ages and is thought to be coincident with impaired phagocytosis and accumulation of lipid deposits (Notomi et al., 2019). RPE morphology is also altered with loss of the normal hexagonal mosaic, suggestive of RPE dysfunction. These features are often seen in age-related macular degeneration (AMD) and RPE-driven inherited retinal diseases. Indeed, we show that conditional loss of *Myrf* impacts the expression of many genes known to be involved in IRD and AMD. We have identified novel targets of *Myrf* that impact RPE cytoskeleton and ultrastructure that may contribute to these structural phenotypes, including *Ermn* and *Upk3b. Ermn,* encodes for Ermin (ERMN), an F-actin binding protein in the ezrin, radixin, moesin superfamily. While it has not been identified as a direct target of *Myrf* in oligodendrocytes (Bujalka et al., 2013), it plays a critical structural role in myelination, and induces cellular protrusions in oligodendrocytes (Brockschnieder et al., 2006). In the RPE, ERMN is specifically expressed in the RPE of adult rat and colocalizes with cytoskeletal markers in the rat and in ARPE19 cells, although its exact role is unknown (Liang et al., 2018). We show that ERMN expression is likewise localized to the apical surface of the RPE and expression is lost in *Myrf* knockouts. Heterozygous *ERMIN* variants have recently been identified in a family with multiple sclerosis, but ocular features such as retinal degeneration or nanophthalmos were not ascertained in this family (Ziaei et al., 2022). *Upk3b* has not been studied in eye development but is a component of the urothelium, which is an epithelial structure composed of extracellular matrix and provides and extensive barrier protecting the urethra from the toxicity of urine (Rudat et al., 2014). It is intriguing to consider its importance in the RPE role as the RPE is critical to forming the blood-retinal barrier.

Based on the expression of *Myrf* in the RPE and the role it plays in RPE development, placing it in the hierarchy of transcription factors is important for understanding its impact in RPE development and disease, and potential as a therapeutic target for RPE degeneration and eye size disorders. The temporal hierarchy of expression of many of the transcription factors known to be involved in RPE development has been established (reviewed in (Amram et al., 2017; Gupta et al., 2023; Martinez-Morales et al., 2004)). Analysis of SCENIC regulon activity in the RPE cluster of our scRNAseq data, predicts comparable activity of the early RPE transcription factors *Pax6, Mitf,* and *Otx2* in wild-type and *Rx>cre Myrf^fl/fl^.* Conditional deletion of *Pax6* in the RPE causes loss of pigmentation by e12.5 but does not affect the ultrastructure of the RPE cells (Raviv et al., 2014). *Pax6* activates *Mitf* and together are important for early activation of genes involved in melanogenesis, as well as specification of the RPE cell fate (Raviv et al., 2014). In our study, we show that RPE are specified in *Rx>cre Myrf^fl/fl^* eyes, they have a similar melanogenesis defects as *Pax6* and *Mitf* knockout mice, and *Pax6* and *Mitf* expression and regulon activity is unchanged. These results support a model in which *Myrf* is downstream of *Pax6* and *Mitf* in the regulation of RPE development and melanogenesis (Figure 10). *Tead1*, a downstream target of the Hippo pathway, was also identified as a regulon active during early RPE development and unchanged during early development in our *Rx>cre Myrf^fl/fl^* mice. Conditional deletion of YAP in the RPE of mice using *Rx>cre* results in transdifferentiation of the RPE to retinal-like cells, and similar transcriptional changes and loss of melanogenesis are observed in yap -/- zebrafish as in our *Rx>cre Myrf^fl/fl^* mice (Kim et al., 2016; Miesfeld et al., 2015). Together, these results suggest that *Myrf* is also downstream of the Hippo signaling pathway. An intriguing finding of our study is that *Sox10* is downstream of *Myrf* in RPE development. *Myrf* and *Sox10* have been shown to be interacting partners in oligodendrocyte differentiation (Aprato et al., 2020; Hornig et al., 2013). *Myrf* has been found to be a direct target of *Sox10* and they work together to switch target genes of oligodendrocyte progenitors to that of differentiation (Aprato et al., 2020). In the eye, Sox10 expression is specifically lost in RPE in our *Rx>cre Myrf^lfl/fl^*, while melanocyte expression remains (Figure 7). Future studies with CUT&RUN or other chromatin immunoprecipitation techniques will help to determine whether these are direct targets and similar regulatory programs act in RPE development as in oligodendrocyte maturation.

**Figure 10.**
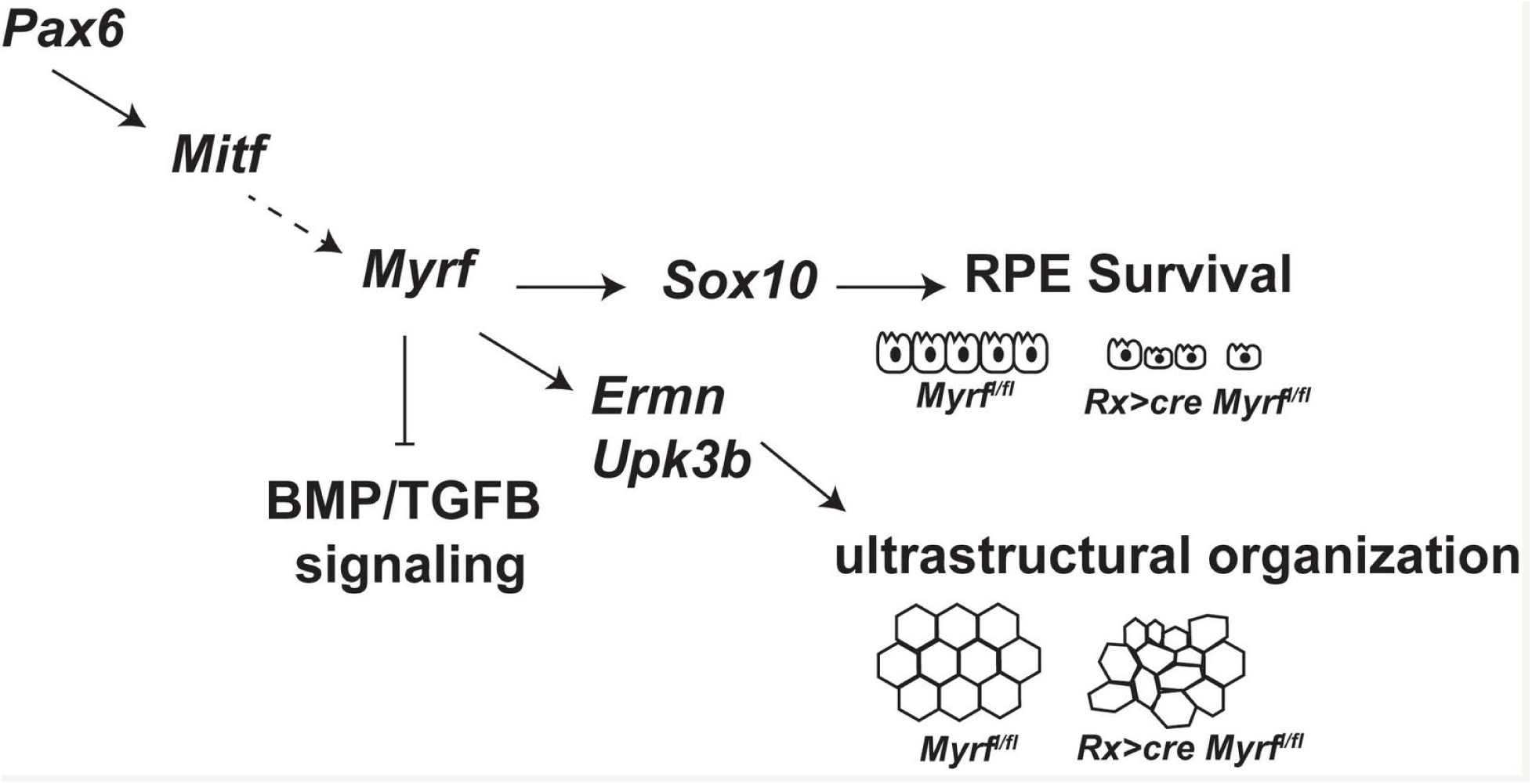
Model. Transcriptional and phenotypic analysis of the *Rx>cre Myrf^fl/fl^* mice suggests *Myrf* is downstream of *Pax6* and *Mitf* in the hierarchy of RPE transcription factor regulation. *Myrf* is important for the activation of *Sox10* which may impact RPE cell survival, *Ermn* and *Upk3b*, which may regulate ultrastructural organization of the RPE, and BMP/TGFB signaling, which may influence eye growth or RPE dysfunction.

We have identified an elevation of TGFβ and BMP signaling in our conditional loss of *Myrf* model, with upregulation of *Tgfb2*, *Bmp2* expression and downstream signaling. These pathways are particularly relevant to retinal and RPE development, RPE function, and eye size. In early stages, TGFβ/BMP signaling from the surface ectoderm has been shown to drive neural retina specification (reviewed in (Amram et al., 2017)). WNT and TGFβ/BMP signaling from the extraocular mesenchyme and surface ectoderm drive RPE cell fate (Steinfeld et al., 2013). In the chick, after removal of the surface ectoderm, application of WNT or BMP-soaked beads is sufficient to activate *Mitf* and initiate RPE specification (Steinfeld et al., 2013). Studies in chick and mice have also shown that BMP2 is a negative regulator of ocular growth (Mai et al., 2022; Zhang et al., 2012). Upregulation of the TGFβ pathway, specifically *Tgfb2*, has been associated with RPE dysfunction (reviewed in (Zhou et al., 2020)). In RPE disease state, TGFβ signaling upregulation induces an inflammatory response, promoting an epithelial to mesenchymal transition (EMT) through both a SMAD dependent and independent pathway (Xu et al., 2009). Tgfb2 induces EMT in cultured human ARPE19 cells (Wu et al., 2018). In our model, condition loss of *Myrf* leads to ultrastructural changes in the RPE, like those observed in Tgfb2 induced EMT. We speculate that the activation of the TGFβ family in *Rx>cre Myrf^fl/fl^* mutants is a response to the induced disease state of the RPE.

Collectively, our study defines a critical role for MYRF in RPE development in melanogenesis, cell structure, and cell survival, places MYRF in the hierarchy of RPE differentiation, and identifies novel candidate genes for RPE-driven disorders. Future studies will be essential to parse out direct targets of *Myrf*, its role in maintaining RPE structure, and the disparate nature of the mouse and human phenotypes.

## MATERIALS AND METHODS

### Mice

*Rxcre* (Swindell et al., 2006) and *Myrf flox* (Emery et al., 2009) mice were housed at the University of Michigan in accordance with guidelines from the Unit for Laboratory Animal Medicine and the Institutional Animal Care & Use Committee. Mice were genotyped from DNA isolated from tail biopsies, as previously described (Garnai et al., 2019).

### RNAscope in situ hybridization

Heads were harvested from embryonic day 14.5 (e14.5) embryos, dissected eyes were harvested from postnatal day 0 (P0) – P21 pups, and optic nerves were harvested from P21 pups. For larger eyes (after P0), a hole was punctured through the cornea to allow penetration of the fixative. Samples were fixed in 4% buffered paraformaldehyde (0.1M NaPO4 pH7.3) for 0.5-4 hours at 22°C, dehydrated through increasing concentrations of ethanol up to 70%. The samples were then embedded in paraffin using the TissueTek VIP Model VIP5A-B1 (Sakura Finetek USA, Inc.) and the Shandon Histocentre 2 Model #64000012 embedding station (Thermo Fisher Scientific) and sectioned at 5-6μm.

For *in situ* hybridization, RNAscope was performed using the RNAscope® Multiplex Fluorescent Detect V2 system (Advanced Cell Diagnostics [ACD], #323110). Briefly, paraffin was removed with two changes of Xylene and then washed in 100% ethanol (ETOH). Sections were treated with the hydrogen peroxide reagent for 10 minutes followed by two washes in distilled water. Target retrieval was performed using boiling 1 X Target Retrieval Reagent for 7 minutes, followed by washing in distilled water and 100% ETOH. After drying the slides, the sections were treated with Protease Plus in prewarmed humidity chamber for 25 minutes at 40°C then washed in distilled water. Prewarmed RNAscope probes were then applied, and slides were incubated in a humidity chamber at 40°C for 2 hours. The probes used were *Mm-Myrf* (ACD, 524061), *Mm-Upk3b* (ACD, 568561), *Mm-Wfikkn2* (ACD, 531321), *Mm-Id3* (ACD, 445881), *Mm-Bmp2-E3* (ACD, 427341), and a negative control probe (ACD, 320871). After hybridization of the probe, sections were washed in 1 X Wash Buffer then incubated at 40°C with Amp1 reagent for 30 minutes, Amp2 reagent for 30 minutes, and Amp3 reagent for 15 minutes, with washes in 1 X Wash Buffer between each step. For signal development, the sections were then incubated in HRP-C1 for 15 minutes at 40°C, washed in 1 X Wash Buffer and then incubated with Cyanine 3, Opal-Tm 570 (Akoya Biosciences, FP1488001KT) fluorophore diluted 1:1500 in TSA plus buffer (ACD, 322809) for 30 minutes at 40°C. Sections were then washed in 1 X Wash Buffer then treated with HRP Blocker for 15 minutes at 40°C, stained with DAPI (Sigma, MBD0015) and mounted in ProLong Gold Antifade (Invitrogen, P36930).

### Single cell RNA sequencing

Eyes were collected from e13.5, e15.5, and P0 litters, the cornea, lens, and optic nerve were removed, and the eyecups were place on ice in 1 X HBSS (Invitrogen, 14175095) while samples were genotyped using a rapid genotyping protocol (Truett et al., 2000). Three *Myrf^fl/fl^* and three *Rx>*c*re Myrf^fl/fl^* samples were pooled for each time point. Eyecups were dissociated into single cells in a Papain solution containing 5mM L Cysteine (Sigma, C7352), 1mM EDTA (Sigma, E4884), 0.6 mM 2-mercaptoethanol (Sigma, M6250), and 1mg/ml Papain (Roche, 10108014001) for 10 minutes at 37°C with trituration every 2 minutes. The dissociation was stopped using Neurobasal Media (Invitrogen, 12348017) with 10% Fetal Bovine Serum (FBS) (Corning, MT35010CV), single cells were collected by centrifugation for 5 minutes at 300 RCF at 4°C. The supernatant was aspirated, and the single cells were resuspended in cold Neurobasal Media with 3% FBS. Single cell libraries were prepared by the University of Michigan Advanced Genomics Core and sequencing was performed using the 10X Genomics Chromium platform using manufacturer’s protocol for 10x Single Cell Expression 3’ and sequenced on the NovaSeq (S4) 300 cycle (Illumina, San Diego, CA). Sequencing outputs were demultiplexed using 10x Genomics Cell Ranger 7.1.0 software and FASTQ files were aligned to the Genome Reference Consortium Mouse Build 38, mm10 genome (Church et al., 2009; Mouse Genome Sequencing et al., 2002; Zheng et al., 2017, https://genomes.ucsc.edu). Seurat/4.1.1 (Butler et al., 2018; Stuart et al., 2019) scRNAseq software was utilized to normalize data using the NormalizeData and FindVariableFeatures commands with nfeatures set to 2000. Poor quality cells were removed using cutoffs of nFeature_RNA < 200 and % mitochondrial RNA >15%. Doublet cells were considered those with nCounts > 2000 and removed from the study. Data from all time points and genotypes was anchored and integrated prior to scaling, principal component analysis and Uniform Manifold Approximation and Projection (UMAP) clustering. The FindAllMarkers command was used to identify and define unique transcripts from each cluster, and cluster identities were defined and assigned using published literature. The FindMarkers command was used to identify differentially expressed genes between clusters and genotypes. Data was displayed in RStudio using the VlnPlot, DotPlot, and FeaturePlot (reduction = “umap”) functions. Gene set enrichment analysis was performed using Gene Ontology Consortium website () using the PANTHER Overrepresentation Test (Release 20200728) and the database GO Ontology database DOI: 10.5281/zenodo.3954044 Released 2020-07-16 (Ashburner et al., 2000; Gene Ontology et al., 2023). Molecular pathway analysis was performed with Differential Nichenetr which analyzes ligand-receptor networks and generates ligand-target matrix (Browaeys et al., 2020; Guilliams et al., 2022), using the RPE cluster as the sender and the RPE cluster as the receiver. Analysis of regulon activity in the RPE cluster, the expression of a given transcription factor compared to the expression of target genes with binding sites, was performed with Single Cell rEgulatory Network Inference and Clustering (SCENIC), using default parameters described in original manuscript (Aibar et al., 2017; Crow et al., 2016). MultiNichenetr and Differential Nichenetr were performed with R packages from the Saeys Lab, using codes provided (Browaeys et al., 2023; Browaeys et al., 2020; Crowell et al., 2020). Pseudotime analysis was performed with the Monocle3 package, using codes provided (Qiu et al., 2017; Trapnell et al., 2014).

### Immunohistochemistry and OTX2/TUNEL staining

Antibody staining was performed using established methods (Garnai et al., 2019), with the antibody conditions can be found in Supplemental Table 3. All antibodies were incubated at 4°C overnight. Nuclei for all samples were stained with DAPI (Sigma, MBD0015) and mounted in ProLong Gold Antifade Mountant (Invitrogen, P36930). RPE flatmounts were done as described previously (Garnai et al., 2019). Co-staining of the rabbit anti-OTX2 (Abcam, ab21990) and apoptotic cells using In Situ Cell Death Detection Kit, TMR red (Roche, 12156792910) was performed on e14.5 paraffin sections. OTX2 immunohistochemistry was performed as described above. Apoptotic cells were identified using the protocol from the In Situ Cell Death Detection Kit, TMR red. Sections pretreated with DNaseI were used as a positive control and sections without the Enzyme Solution were used as a negative control. Nuclei were stained with DAPI (Sigma, MBD0015) and mounted in ProLong Gold Antifade Mountant (Invitrogen, P36930). OTX2 positive, TUNEL positive, and DAPI positive cells were counted using Fiji, ImageJ software. A minimum of two slides were counted per sample. For the OTX2 high counts, the data is displayed as the total number of the highest intensity OTX2 stained cells per total number of DAPI cells in the region counted. For the OTX2 low counts, the data is displayed as the total number of less intensely stained OTX2 cells per the total number of DAPI cells in the region counted. For the RPE apoptosis counts, the data is displayed as the total number of TUNEL positive cells per the total number of RPE cells, marked by OTX2. Each point on the graph represents an individual sample. Statistical analysis was performed using the unpaired Student’s T-Test.

### Electron microscopy

Eyes were collected from P21 mice and fixed in 2% paraformaldehyde 2% glutaraldehyde in 100 mM cacodylate buffer overnight. The tissue was then rinsed in PBS, post-fixed in osmium tetroxide for 1 hour. Tissues with rinsed 3X in PBS followed by graded alcohol dehydration, and 2X rinses in propylene oxide for 10 minutes each. Eyecups were infiltrated with Epon embedding media by incubation in with propylene oxide:Epon mixes 3:1 then 1:1 then 3:1 each for 1 hour and subsequently 100% Epon overnight while gently agitating the tissue. The eyecups were then embedded in a beam capsule with 100% Epon and incubated at 60°C overnight to cure the resin. The eyecups were then cut into semithin (500 nm) section using a Ultracut E ultramicrotome (Leica Biosystems). Sections were stained with toluidine blue to identify key target areas. The block was then trimmed into 1 mm section for further EM processing. Ultrathin sections (70-90 Å) were collected with a diATOME diamond knife, placed on cooper grids, and counterstained with Uranyless (samarium/gadolinium triacetate) and lead citrate and imaged on the JEOL JSM 1400 Plus transmission electron microscope (JEOL,Peabody,MA) at the Microscopy and Image Analysis Laboratory. Melanosomes were identified within the high-resolution image by inspection and cells were manually segmented to define the cell area. The number of melanosomes per mm^2^ were calculated. Comparison among genotypes was done by student’s T-test.

## ACKNOWLEDGEMENTS

The authors are grateful to the Advanced Genomics Core for assistance with single cell RNA sequencing; to Athera Yakoo for technical assistance; to Ben Emery for Myrf floxed mice and helpful advice; to Elior Peles for the ERMN antibody; to Sally Camper, Jun Li, and David Zacks, Brian Brooks, Kapil Bharti, and Robert Hufnagel for helpful discussions.

## DATA AVAILABILITY

Single cell RNA sequencing data will be deposited in GEO (Accession number: ******). Primary data is provided for all of the experiments. All additional relevant data can be found within the article and its supplementary information and primary data are available upon request from the corresponding author.

## WEB RESOURCES

Eye Integration: https://eyeintegration.nei.nih.gov/

UCSC Genome Browser: https://genomes.ucsc.edu

